# MSH2 and MSH6 as size dependent cellular determinants for prime editing in human embryonic stem cells

**DOI:** 10.1101/2022.08.17.504216

**Authors:** Ju-Chan Park, Yunjeong Kim, Jun Hee Han, Dayeon Kim, Jumee Kim, Hyeon-Ki Jang, Sangsu Bae, Hyuk-Jin Cha

## Abstract

Potential applications of precise genome editing in human pluripotent stem cells (hPSCs), not only for isogenic disease modeling but also for *ex vivo* stem cell therapy, have urged the application of diverse genome editing tools in hPSCs. However, unlike differentiated somatic cells, the unique cellular properties of hPSCs (e.g., high susceptibility to DNA damage and active DNA repair) largely determine the overall efficiency of editing tools. Considering high demand of prime editors (PE), mostly due to its broad editing coverage compared to base editors, it is important to characterize the key molecular determinants of PE efficiency in hPSCs. Herein, we showed that MSH2 and MSH6, two main components of the MutSα complex of mismatch repair (MMR), are highly expressed in hPSCs and determine PE efficiency in an ‘editing size’-dependent manner. Importantly, loss of *MSH2,* which disrupts both MutSα and MutSβ complexes, was found to dramatically improve the efficiency of PE from one base to 10 bases, up to 50 folds. In contrast, genetic perturbation of *MSH6,* which solely abrogates MutSα activity, marginally improved the editing efficiency up to 3 base pairs. The size dependent effect of MSH2 or MSH6 on prime editing in hPSCs not only implies MMR is a major determinant of PE efficiency in hPSCs but also highlights the distinct roles of MutSα and MutSβ in the outcome of genome editing.

## Introduction

Soon after the establishment of human induced pluripotent stem cells (iPSCs) [1], iPSCs from patients with genetic diseases have been demonstrated as a powerful tool to mimic cellular phenotypes of genetic origin (i.e., disease modeling) [2, 3]. With continuous advancements in the clustered regularly interspaced short palindromic repeats (CRISPR)/Cas9 system technology, patient-derived iPSCs have been recognized not only for their potential for isogenic disease modeling [4, 5], but also as cell resources for *ex vivo* autologous cell therapy following pathogenic gene correction [6]. As the majority of pathogenic genetic variants found in humans are point mutations (58%) and deletions (25%) [7], precise genome editing to enable permanent correction of these genetic variations becomes crucial for clinical applications. To this end, various Cas9-based genome editing technologies, including homology directed repair (HDR) based knock-in (KI) [8], base editors (BEs) [9] and prime editor (PE) [10], have been applied in human pluripotent stem cells (hPSCs) and optimized for improving efficiency [11, 12].

Unlike introducing random insertion/deletion (indel) for knock-out (KO), precise genome editing technologies utilize distinct editing strategies to modify genome. When supplemented with donor DNA, the Cas9 enzyme conducts precise genome editing via homologous recombination [13]. However, only a small population achieves desirable KI while the majority of cells gain random indel mutations by non-homologous end joining (NHEJ) pathway, an error prone DNA double-strand break (DSB)-repair pathway [13]. In addition, endonuclease activity of Cas9 also cause large deletions and even complex chromosomal rearrangements, resulting in unexpected DNA changes and pathogenic consequences [14, 15]. In particular, patient derived iPSCs of which pathogenic mutations are corrected by HDR based KI, are revealed to be hemizygous due to large on-target defects [16]. Alternatively, BEs achieve base transitions without DSB, by deaminase conjugated to nickase Cas9 (nCas9) or catalytically dead Cas9 (dCas9) [17, 18]. Thus, higher single base substitution efficiency than conventional HDR-based KI [9] and no large deletion [19] renders BEs beneficial and secure tools for gene correction of pathogenic mutations of hPSCs. However, the limited scope of base substitution (only transition and C-to-G transversion) and target availability within editing activity window limits the coverage of pathogenic genetic variations [9]. Considering this, PE enables a highly flexible genome editing such as targeted introduction of indels and base substitutions through the synthesis of a DNA strand into a target site via reverse transcriptase (RT) and an engineered PE guide RNA (pegRNA) [20]. Due to the wider coverage of PE (e.g., transition/transversion point mutation and short insertion/deletion) compared to BE, extensive studies have been performed to improve PE editing efficiency [21–23].

Embryonic stem cells (ESCs), where all somatic cells in body originate during development, have evolved molecular mechanisms to maintain genome integrity, which rely on the high susceptibility to DNA damages and active DNA repair pathways [24–26]. These unique characteristics of ESCs (also in hESCs and iPSCs) serve as major determinants for the outcome of genome editing tools. For instance, relatively low Cas9-mediated genome editing efficiency in hPSCs [27] was recently explained by high p53-dependent cell death upon DSB by Cas9 [28]. In addition, lower overall efficiency of cytosine base editor (CBE) than that of adenine base editor (ABE) in hPSCs results from the active base excision repair (BER) by elevated expression of the uracil DNA glycosylase (UNG) [29]. Despite two recent pioneering studies to examine PE in hPSCs [10, 30], little is known about the unique cellular properties of hPSCs in outcomes of PE.

DNA mismatch repair (MMR) is a highly conserved biological pathway that plays a key role in the maintenance of replication fidelity by a series of processes, namely mismatch recognition, mismatch removal, and synthesis of repair DNA [31]. DNA mismatch is recognized by mismatch repair complexes MutSα (MSH2-MSH6 subunits, the most abundant components in MMR) and MutSß (MSH2-MSH3 subunits) [31]. The MutSα initiates base mispairs (e.g., G-U mispair by cytosine deamination) as well as small insertion/deletion loops (IDLs) of one or two extrahelical nucleotides to cover the repair of most replication error [32, 33]. In contrast, MutSβ governs the repair of most large IDLs [31].

Herein, we demonstrated that *MSH2* (MutS homology 2) and *MSH6* (MutS homology 6), key components in MMR, are highly expressed in hPSCs and serve as cellular determinants for PE efficiency according to editing sizes. Consistent with previous studies [21, 22], knockout of *MSH2*, which leads to the functional inactivation of MutSα and MutSβ, dramatically improved the efficiency of PE on a single base pair to 10 base pairs in hPSCs. In contrast, genetic perturbation of *MSH6,* which only impedes the actions of MutSα on MMR, improved the efficiency of PE on up to 3 base pairs. Considering the critical roles of MMR in the regulation of genome integrity of actively proliferating cells (i.e., hPSCs), the transient modulation of MutSα or MutSβ components, depending on the desired editing size, could be considered as a secure strategy to improve the prime editing in hPSCs.

## Material and Methods

### Cell line and culture and transfection

Human embryonic stem cells (WA09, WiCell Research Institute) were cultured on Magrigel (BD Biosciences) coated cell culture dish in StemMACS media (Miltenyi-Biotec) added with 50 μg/ml Gentamicin (Gibco). At 70 ~ 80 % of confluency, cells were rinsed with DPBS (Gibco) and exposed to Accutase (BD Biosciences) for detachment. Detached cells were washed with DMEM/F-12 (Gibco) media and plated to Matrigel-coated plate fed with StemMACS media added with 50 μg/ml Gentamicin (Gibco) and 10 μM of Y27632 (Gibco). Cells were detached with Accutase solution (561527, BD Biosciences) and washed with Opti-MEM (31985070, Gibco) for three times and diluted to concentration of 1 x 106 cell in 100 μl of Opti-MEM for transfection. 2 μg of PE2 of Cas9 vector and 3 μg of sgRNA or pegRNA vector was added to cell mixture. Electroporation was performed by NEPA-21 with 175 V of poring pulse and 2.5 ms of pulse length.

### Targeted deep sequencing

For analysis, genomic DNA (gDNA) of each sample was extracted by NucleoSpin Tissue kit (MN) following the supplier’s instructions. To generate sequencing libraries, target sites were amplified using a TOYOBO KOD Multi & Epi PCR kit. As previously disclosed, these libraries were sequenced utilizing MiniSeq and an Illumina TruSeq HT Dual Index system. Using an Illumina MiniSeq technology, equal amounts of PCR amplicons were submitted to paired-end read sequencing. Paired-end reads were analyzed by comparing mutant and WT sequences using BE-analyzer after MiniSeq.

### RT-qPCR

Easy-BLUE^TM^ RNA isolation kit (iNtRON Biotechnology) was used for total RNA extraction following the supplier’s instructions. cDNA was synthesized by adding 2 μl of PrimeScriptTM RT reagent kit (TaKaRa) to 500μg of RNA samples in 8 μl of distilled water (DW) and reacted for 15 min in 37 °C. Synthesized cDNA was diluted in 200 μl of DW and 9 μl of diluted cDNA was added to 96 well qPCR plate. 1 μl of qPCR primer and 10 μl of SYBR^®^ Green PCR reagents (Life Technologies) were added to each well. qPCR was performed by Light Cycler-480^®^II (Loche).

### Gene Set Enrichment Analysis (GSEA) and KEGG-Pathway analysis

Four selected gene expression profiling by RNA microarray (GSE9709, GSE2248, GSE16963, GSE42445) were downloaded from the Gene Expression Omnibus (GEO) database. Rank for differently expressed RNA/genes (DEGs) between hESCs and differentiated cell were yielded by GEO2R(https://www.ncbi.nlm.nih.gov/geo/geo2r/). Using the gene list as input, GSEA analysis was performed using the KEGG_MISMATCH_REPAIR genesets from MSigDB via the R package “fgsea”, and KEGG-pathway analysis was performed by the R package “pathview (https://doi.org/10.1093/bioinformatics/btt285)”. Significantly upregulated or downregulated GO terms were selected with nominal p-value < 0.05 and a Normalized enrichment score (NES)|>1.5.

### Statistical analysis

The quantitative data are expressed as the mean values ± standard deviation (SD). Student’s unpaired t-tests was performed to analyze the statistical significance of each response variable using the PRISM. p values less than 0.05 were considered statistically significant (*, p < 0.05, **, p <0.01, ***, p <0.001 and ****, p <0.0001)

## Results

### High enrichment of MMR pathway in hPSCs

Editing outcome of PE was associated with the MMR activity (Fig. 1A) [ref]. nCas9 of PE produces single strand break (SSB) near the PAM sequence (a) and the RT reaction next synthesizes edited DNA (3’ flap) based on the RT template sequence of the pegRNA (b). The 3’ flap is annealed to the non-edited strand, producing 5’ flap through an equilibrium process (c). Following removal of the liberated 5’flap by the DNA damage repair machinery (d), the intended edit is incorporated into the target DNA sequence (e). The 5’ flap excised intermediate product is susceptible to MMR that results in the removal of newly synthesized sequence (f) (Fig. 1A). This implies that MMR activity, varied in cell type specific manner [34], determines PE efficiency. Consistently, recent genome wide screening reveals that repression of major components of MMR including MSH2, MSH6 or MLH1, enhances efficiency and fidelity of PE to lead to the improvement of PE version (i.e., PE4 and PE5) through the transient modulation of MMR activity [22].

**Figure 1.**
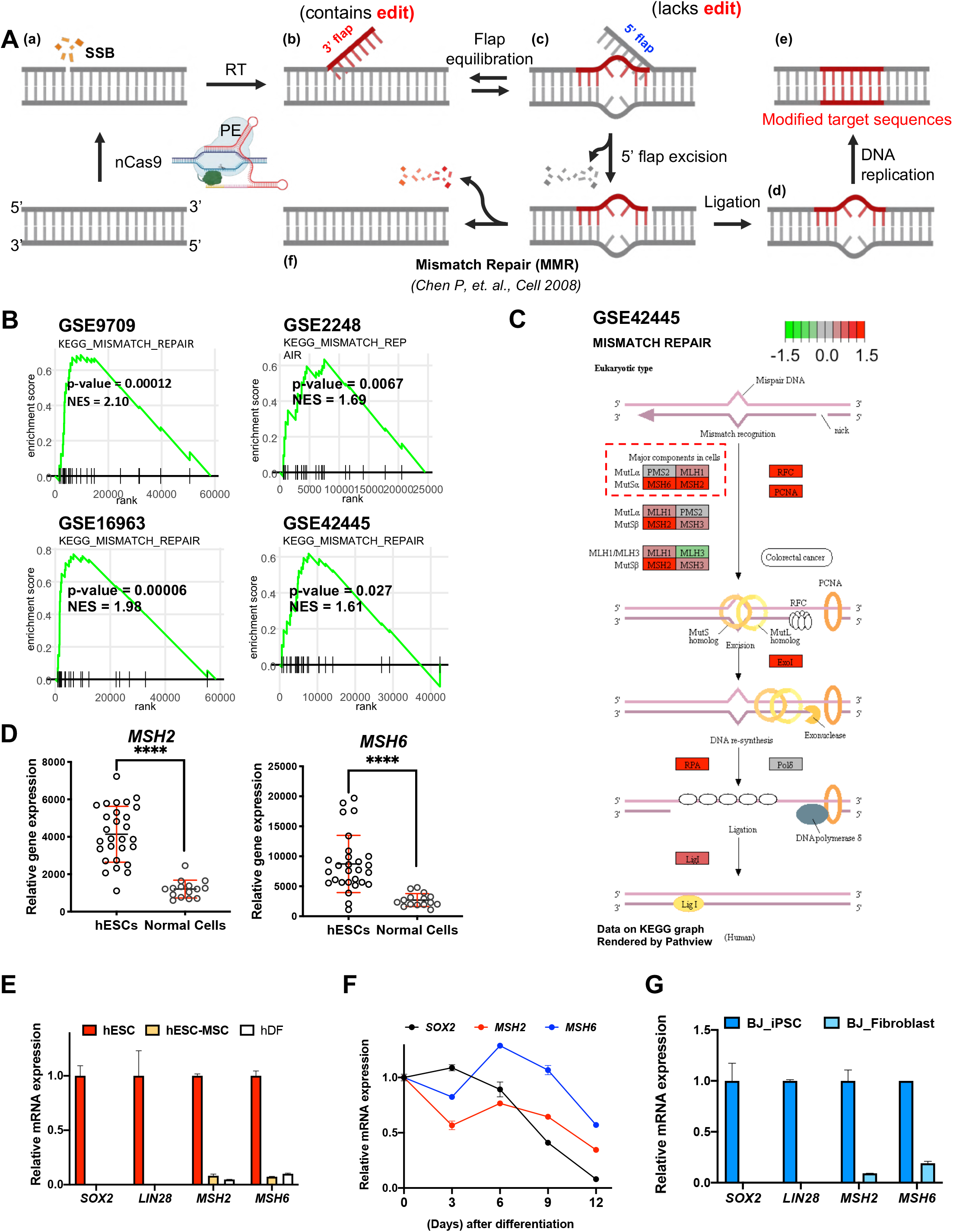
High enriched mismatch repair pathway and high expression of MSH2 and MSH6 in hPSCs. (A) Scheme of prime editing process (B) Gene set enrichment analysis for gene ontology of miamatch repair pathway from indicated gene sets (C) KEGG pathway analysis of eukaryotic mismatch repair from indicated gene set, highly expressed genes are colored in red while lowly expressed genes are colored in green. (D) Expression of MSH2 and MSH6 in several hESCs cell lines from NextBio portal, Normal cells were analyzed as controls. Bars represent mean values, and error bars represent the S.D. ****P < 0.0001, comparing with the hESCs by unpaired t-test. (E) mRNA expression of SOX2, LIN28, MSH2 and MSH6 of hESC-derived MSCs (H9-MSCs), hESCs (H9-ESCs) and human dermal fibroblasts (hDF). Bars represent mean values, and error bars represent the S.D. (n = 2) (F) mRNA expression of SOX2, MSH2 and MSH6 from H9-hESCs at indicative days after differentiation, Dots represent mean values, and error bars represent the S.D. (n = 2) (G) mRNA expression of SOX2, LIN28, MSH2 and MSH6 of human fibroblast (BJ-Fibroblast) and BJ-fibroblast induced iPSCs (BJ-iPSCs), Bars represent mean values, and error bars represent the S.D. (n = 2)

As previously demonstrated for CBE, in which the efficiency is determined by active BER [29], we hypothesized that unique cellular characteristics of hPSCs, in which MMR is highly activated, may affect PE efficiency. As expected, multiple gene sets associated with MMR were highly enriched in undifferentiated hPSCs compared to differentiated counterparts (Fig. 1B). To illustrate key components of high MMR activity in hPSCs, upregulated genes of undifferentiated hPSCs were next depicted in the KEGG pathway map. Consistent in all datasets (i.e., GSE9709, GSE2248, GSE16963 and GSE42445) (Fig. S1A), *MSH2* and *MSH6* were highly expressed in undifferentiated hPSCs (Figs. 1C, S1B-D) among known cellular determinants of PE efficiency (e.g., *MLH1, PMS2, MSH2,* and *MSH6)* [22]. The high expression levels of *MSH2* and *MSH6* of total 25 different hESCs lines compared to 14 normal cells, were again validated by nextbio dataset (http://nextbio.com) (see Table S1 for full list of cell information).

### Distinctive expression levels of *MSH2* and *MSH6* genes in hPSCs

As predicted by hPSCs transcriptome datasets (Fig. 1), *MSH2* and *MSH6* gene expression was highly expressed in hESCs (of which pluripotency was determined by expression levels of *SOX2* and *LIN28)* compared to that of human mesenchymal stem cells derived from hESCs (hESC-MSCs) [35] and human dermal fibroblasts (hDFs) (Fig. 1E). In addition, mRNA levels of *MSH2* and *MSH6* were markedly repressed during spontaneous differentiation of hESCs (Fig. 1F).. The unique expression patterns of MSH2 and MSH6 were also validated in iPSCs (e.g., BJ-iPSCs) compared to BJ fibroblast, a parent somatic cell of iPSCs [36] (Fig. 1G). Considering the correlation of protein expression level of MMR components with overall MMR activities in cell lines [34], high expression of MSH2 and MSH6 in hPSCs implied their MMR activity.

### Genetic perturbation of *MSH2* or *MSH6* in hESCs with CRISPR/Cas9

In order to examine the effects of a higher expression of *MSH2* or *MSH6* on PE efficiency in hPSCs, *MSH2* or *MSH6* was genetically perturbed by CRISPR/Cas9 (hereafter Cas9). In order to improve Cas9-mediated editing outcome in hPSCs [28], un-edited hPSCs after Cas9 treatment were selectively eliminated by treatment with YM155, a survivin inhibitor used to selectively kill undifferentiated hPSCs [37] through SLC35F2, which is highly expressed in undifferentiated hPSCs [12]. The enriched selection procedure was performed as previously demonstrated [12], using YM155 for efficient knockout of gene of interest (GOI) (Fig. 2A). Through the enrichment of edited hPSCs by this approach, single clone of MSH2 homozygous knockout (MSH2^-/-^: M2^-/-^, Fig. 2B) and MSH6 heterozygous knockout (MSH6^-/+^: M6^-/+^, Fig. 2C) along with SLC35F2 homozygous knockout (SLC35F2 KO), were established (Fig. S2A). SLC35F2 KO clone with intact MSH2 or MSH6 were used as KO controls (Mock KO: Cont) (Figs. 2B and 2C). Immunoblotting analysis to determine MSH2 protein levels clearly confirmed the successful establishment of MSH2 KO hESCs (Fig. 2D). Notably, while MSH6 protein level was markedly diminished in M6^-/+^ clones, as expected, a drastic reduction of MSH6 protein was also manifested in M2^-/-^ clones (Fig. 2D), despite compensatory transcription of *MSH6* occurring in the absence of MSH2 (Fig. S2B). This finding was consistent with data from previous studies suggesting that MSH6 is destabilized in the absence of MSH2 [38, 39]. The expression levels of typical pluripotency genes (i.e., *POU5F1, SOX2* and *NANOG)* (Fig. 2E) and proteins (i.e., OCT4 and NANOG) (Fig. 2F) revealed that depletion of *MSH2* or *MSH6* gave only a marginal effect on the pluripotency of hESCs. In fact, the typical colonial morphology and alkaline phosphatase activity demonstrated that these hESCs clones (i.e., Cont, M6^-/+^ and M2^-/-^ hESCs) maintained pluripotency (Fig. 2G).

**Figure 2.**
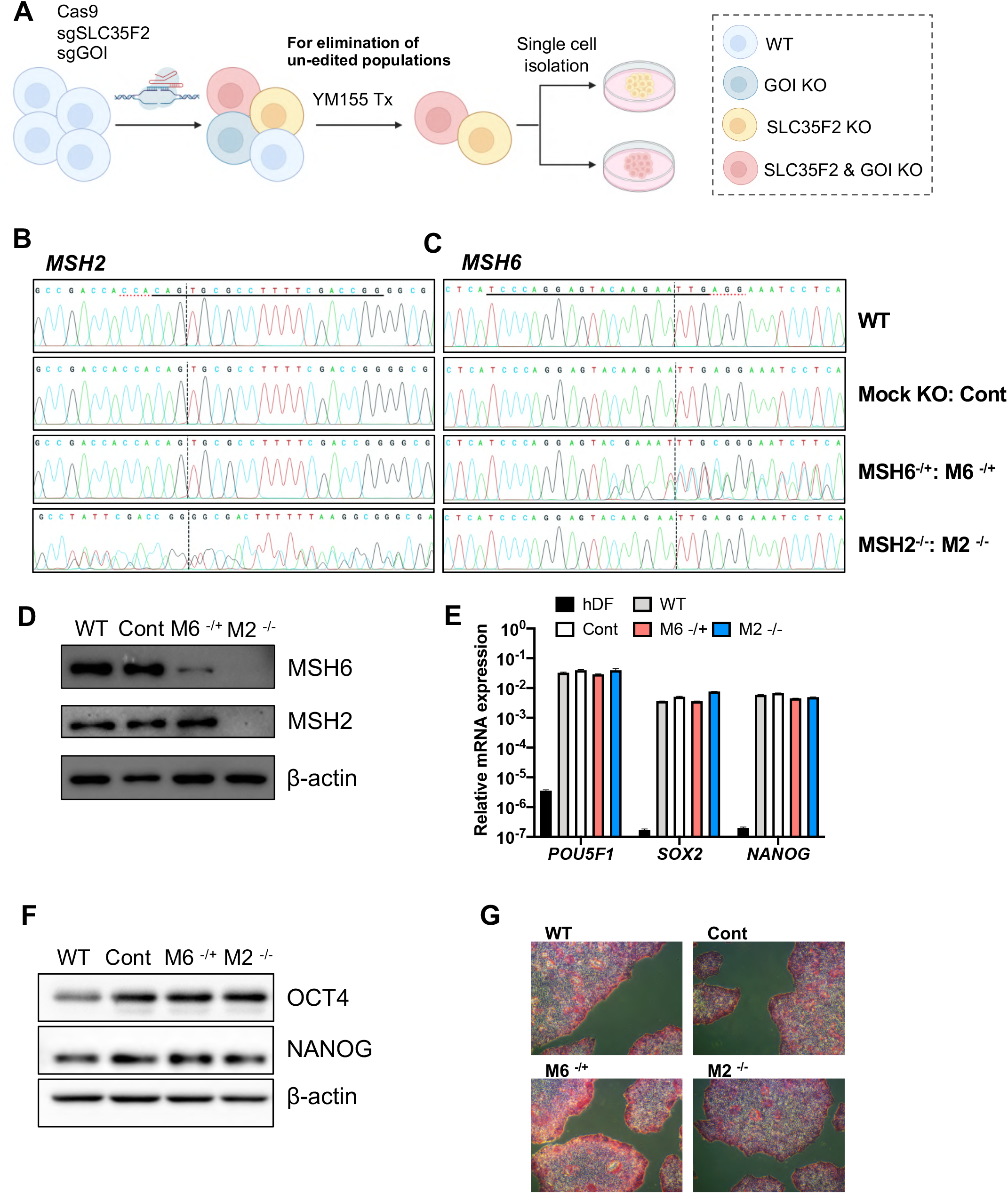
Genetic perturbation of MSH2 or MSH6 in hESCs with CRISPR/Cas9. (A) Scheme of YM155-enriched selection method for GOI KO hESCs establishment (B) Sanger sequencing result of MSH2 in WT, SLC35F2 only KO (Cont), MSH6 heterozygous KO (M6 -/+) and MSH2 homozygous KO (M2 -/-) hESCs (C) Sanger sequencing result of MSH6 in WT, SLC35F2 only KO (Cont), MSH6 heterozygous KO (M6 -/+) and MSH2 homozygous KO (M2 -/-) hESCs (D) Immunoblotting for MSH2 and MSH6 in WT, SLC35F2 only KO (Cont), MSH6 heterozygous KO (M6 -/+) and MSH2 homozygous KO (M2 -/-) hESCs (E) mRNA expression of POU5F1, SOX2 and NANOG in WT, SLC35F2 only KO (Cont), MSH6 heterozygous KO (M6 -/+), MSH2 homozygous KO (M2 -/-) hESCs and human dermal fibroblast (hDF) (F) Immunoblotting for OCT4 and NANOG in WT, SLC35F2 only KO (Cont), MSH6 heterozygous KO (M6 -/+) and MSH2 homozygous KO (M2 -/-) hESCs (G) Alkaline phosphatase activity assay of WT, SLC35F2 only KO (Cont), MSH6 heterozygous KO (M6 -/+) and MSH2 homozygous KO (M2 -/-) hESCs

### Discrimination of editing size of PE by MSH2 or MSH6

As previously mentioned, MSH2 serves as a key component for both MutSα and MutSβ by forming heterodimer with MSH6 and MSH3 to govern size dependent DNA errors, respectively (Fig. 3A). According to the protein levels of MSH2 and MSH6 in M2^-/-^ or M6^-/+^ hESCs (Figs. 2D and E), M2^-/-^ or M6^-/+^ hESCs established in this study, were examined to determine the effect of lacking both MutSα and MutSβ (i.e., M2^-/-^ clone) or only MutSα (with intact MutSβ, i.e., M6^-/+^ clone) on PE efficiency. Considering the distinct roles of MutSα and MutSβ in the recognition of base mispair and IDLs (with different size) [31–33], the different outcomes of PE efficiency were predicted (Fig. 3B). To test this hypothesis, we first determined the PE editing efficiency for one base transversion (i.e., T to A), one base pair insertion/deletion (i.e., A ins and T del) and three base pair insertion (i.e., CTT or GTA ins), of which mutations are mostly repaired by MutSα-dependent MMR. As expected, the M6^-/+^ clone, displaying a partially disrupted MutSα due to diminished levels of the MSH6 protein (Fig. 2E), showed a significant improvement in the editing efficiency for one base (Fig. 3C) and three base pairs insertion (Fig. 3D), while drastic improvement of one base (Fig. 3C) and three base pairs (Fig. 3D) was manifested in the M2^-/-^ clone, lacking both MutSα and MutSβ. In contrast, the PE-mediated insertion of larger IDLs, mostly recognized in MutSβ-dependent manner, was not significantly improved in the M6^-/+^ clone, in contrast to the significant improvement of efficiency in the M2^-/-^ clone (Fig. 3E). However, in the case of the insertion/deletion of over 15 nucleotides, which MutSα nor MutSβ are incapable of recognizing [40], the editing efficiency remained comparable following the loss of MSH2 or MSH6 (Fig. 3F). These results suggest that an increased MMR activity in hPSCs, due to the expression of MSH6 or MSH6, determines the editing efficiency of PE in an editing sizedependent manner.

**Figure 3.**
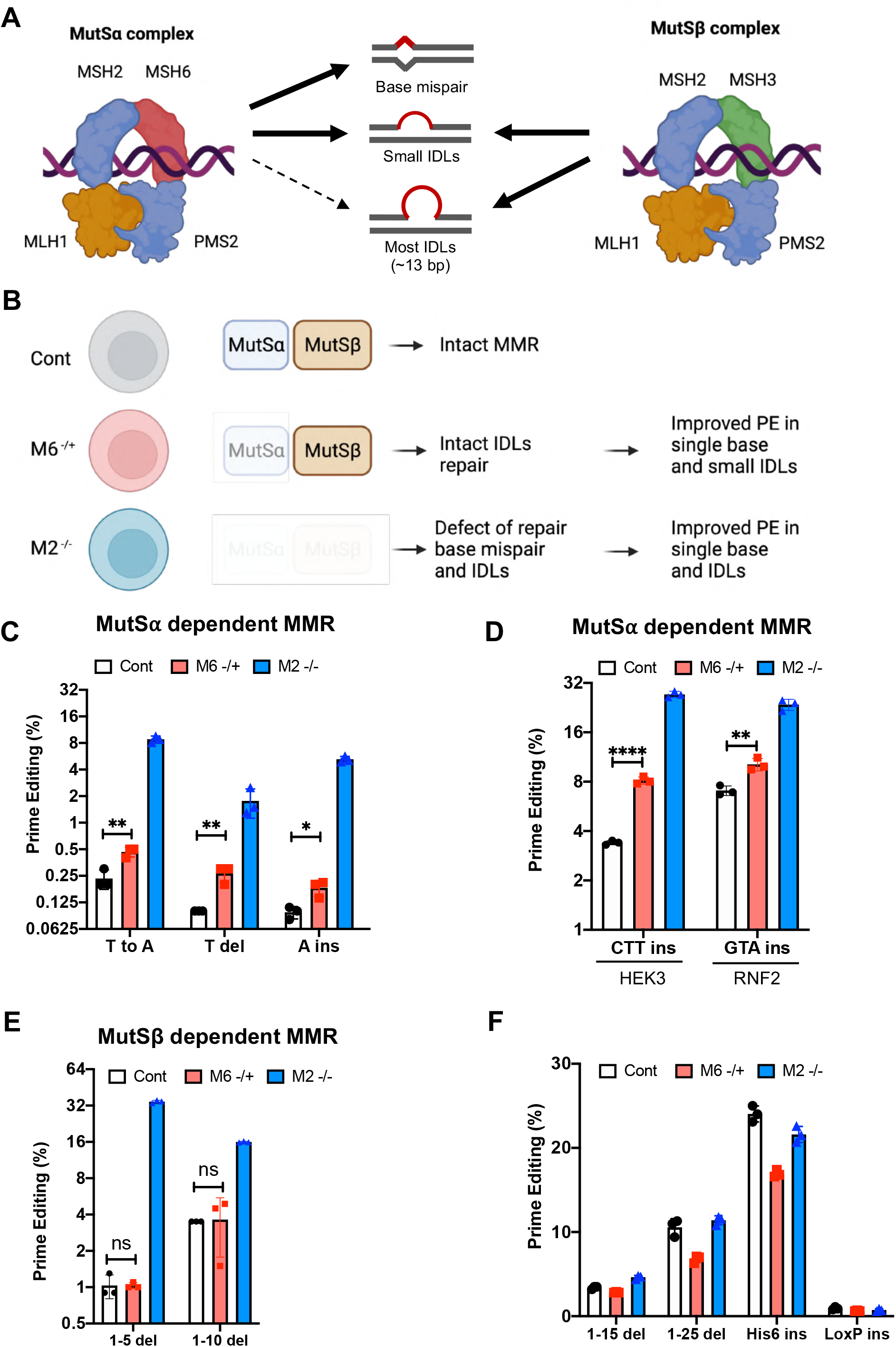
Discrimination of editing size of PE by MSH2 or MSH6. (A) Scheme of mismatch recognition by MutSα and MutSβ complex (B) Scheme of MutSα and MutSβ status in SLC35F2 only KO (Cont), MSH6 heterozygous KO (M6 -/+) and MSH2 homozygous KO (M2 -/-) hESCs (C) Prime editing efficiency inducing T to A, T deletion (T del) and A insertion (A ins) at HEK3 site in SLC35F2 only KO (Cont), MSH6 heterozygous KO (M6 -/+) and MSH2 homozygous KO (M2 -/-) hESCs (D) Prime editing efficiency inducing CTT insertion (CTT ins) at HEK3 and GTA insertion (GTA ins) at RNF2 site in SLC35F2 only KO (Cont), MSH6 heterozygous KO (M6 -/+) and MSH2 homozygous KO (M2 -/-) hESCs (E) Prime editing efficiency inducing 1-5 deletion (1-5 del) and 1-10 deletion (1-10 del) at HEK3 site in SLC35F2 only KO (Cont), MSH6 heterozygous KO (M6 -/+) and MSH2 homozygous KO (M2 -/-) hESCs (F) Prime editing efficiency inducing 1-15 deletion (1-15 del), 1-25 deletion (1-25 del), His 6x tag insertion (His6 ins) and LoxP insertion (LoxP ins) at HEK3 site in SLC35F2 only KO (Cont), MSH6 heterozygous KO (M6 -/+) and MSH2 homozygous KO (M2 -/-) hESCs

## Discussion

The overall PE editing efficiency in hPSCs with a standardized protocol (i.e., PE2 editing), was around 1-3%, which would urge the laborious clonal selection for either disease modeling or gene correction of hPSCs. Considering this, recently developed PE4 or PE5, conjugated by dominant negative MLH1 (dnMLH1) to transiently disrupt both MutSα and MutSβ [22], would be applicable to improve the efficiency of PE in hPSCs. However, the editing outcomes of newly developed editing tools are frequently monitored in cancer cell lines. Therefore, and contrastingly, precise base editing in hPSCs should avoid inadvertent mutations, which may cause unexpected outcomes in not only disease modeling but also in gene correction approaches used for *ex vivo* cell therapy. It is noteworthy that loss of *MLH1* (or *MSH2)* causes a higher frequency of mutations compared to other the loss of other MMR components, such as *MSH3* or *MSH6* [41]. Accordingly, the application of PE4 or PE5, newly developed PE tools based on transient suppression of MLH1, a key MMR protein for both MutSα and MutSβ, may need to be closely examined, whether any unintended mutations occur by simultaneous inhibition of MutSα and MutSβ in rapidly proliferating hPSCs with lack of cell cycle checkpoints [26]. As demonstrated in this study, MutSα or MutSβ serves size dependent cellular determinant of PE outcomes. Thus, instead of disrupting MLH1 or MSH2, leading to the dual inhibition of both MutSα and MutSβ, transient inhibition of MutSα (by suppression of MSH6) or MutSβ (by suppression of MSH3), according to desired editing size respectively, would most likely achieve a safer PE editing of hPSCs.

Additionally, we noted that the basal MMR activity in human cell lines varied, with SW480, CaCo2, HEL, HeLa, MRC and WI38 cell lines being MMR-proficient and HCT116, LoVo, DLD1, HCT15 and SW48 cell lines being MMR-deficient [38]. Accordingly, the editing efficiency of PE4 or PE5, designed to modulate MMR activity, would work in a cell-type specific manner, underlying the need to consider cellular context when selecting appropriate editing tools.

## Supporting information

Supplemental Informations

## Data availability

Source Data are available from the corresponding authors upon request.

---

**Figure. S1.**
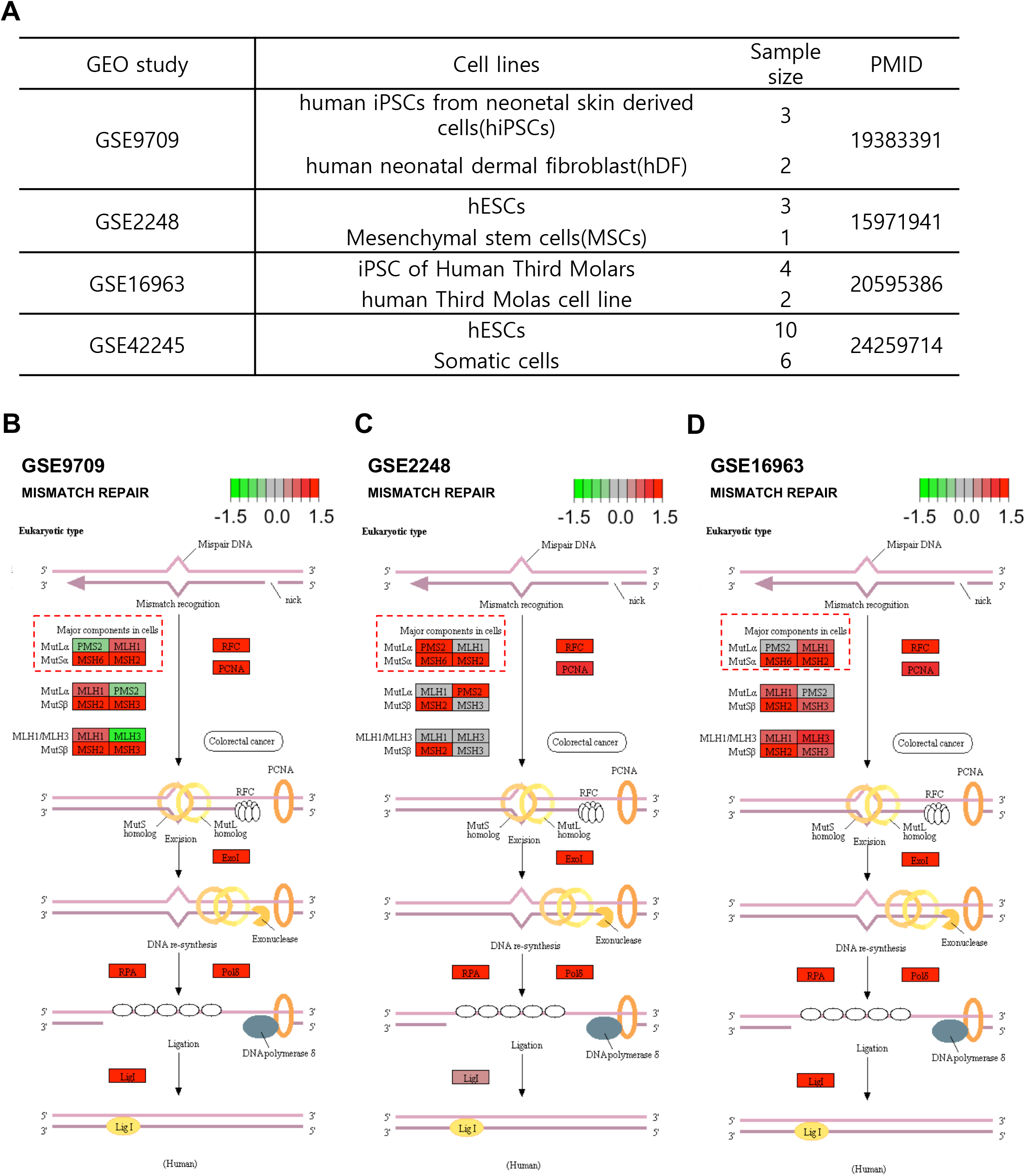

**Figure. S2.**
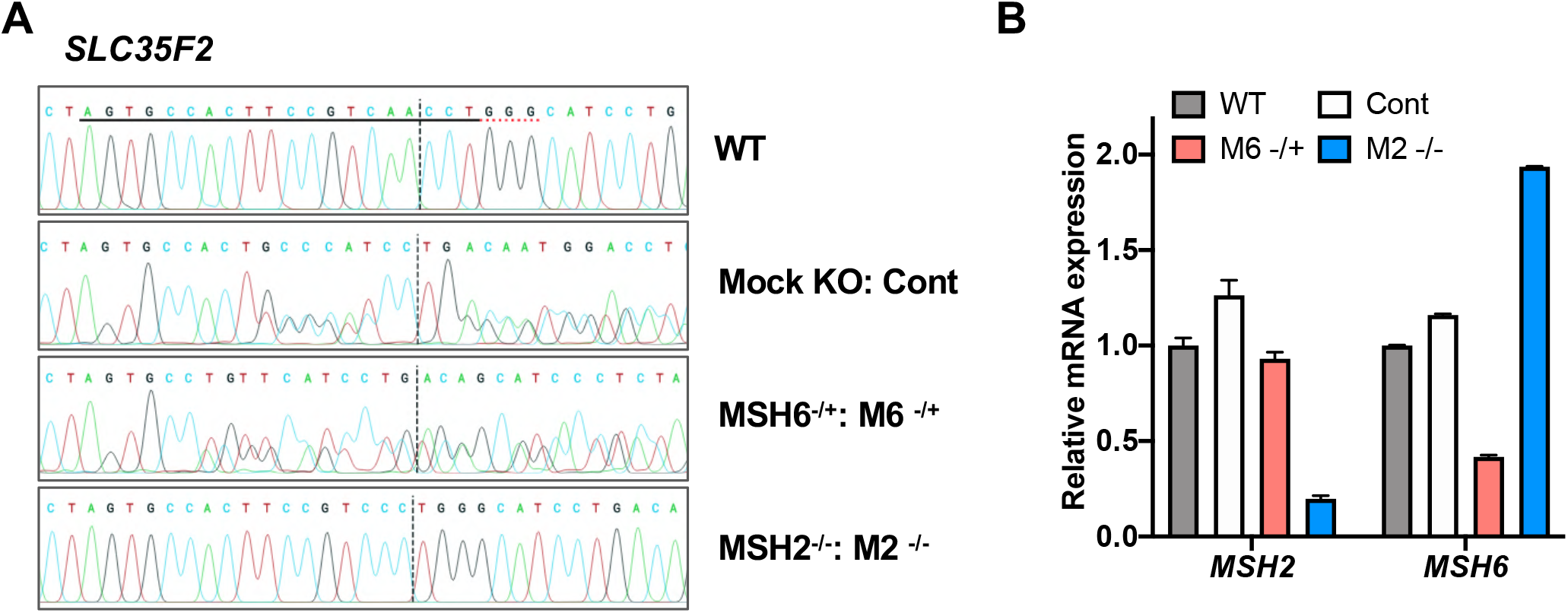

