## Supplemental Informations for "MSH2 and MSH6 as size dependent cellular determinants for prime editing in human embryonic stem cells": 220816_Supplementary Materials_BioRxiv.pdf

**Discrimination of editing size by MSH2 or MSH6 for prime editing efficiency in human pluripotent stem cells**

This PDF file includes:

Experimental Section

Supplemental Figures

Supplemental Figure legends

Supplementary Table 1

**Sequence information**

**2. qPCR primer information**

| Gene | qPCR primer sequence (5' to 3') | |
|---|---|---|
| POU5F1 | F | TGTACTCCTCGGTCCCTTTC |
| | R | TCCAGGTTTTCTTTCCCTAGC |
| NANOG | F | AAATTGGTGATGAAGATGTATTCG |
| | R | GCAAAACAGAGCCAAAAACG |
| SOX2 | F | TTCACATGTCCCAGCACTACCAGA |
| | R | TCACATGTGTGAGAGGGGCAGTGTGC |
| LIN28A | F | CACGGTGCGGGCATCTG |
| | R | CCTTCCATGTGCAGCTTACTC |
| MSH2_1 | F | GCTTCGTGCGCTTCTTTCAG |
| | R | GTATAGAAGTCGCCCCGGTC |
| MSH6_1 | F | CTGAATCCCAAGCCCACGTT |
| | R | CCTCCTCCTGTGCTCATCTC |
| MLH1 | F | ATGCCCACCAGATGGTTCGT |
| | R | CCCTTTGTTGTATCCCCCTCC |
| PMS2 | F | AGAGAACAAGCCTCACAGCC |
| | R | GAACCCCTCAGAATCCACGG |
| 18s rRNA | F | GTAACCCGTTGAACCCCATT |
| | R | CCATCCAATCGGTAGTAGCG |

**2. sgRNA sequence information**

| Gene | sgRNA sequence (w/ PAM) |
|---|---|
| SLC35F2 | AGTGCCACTTCCGTCAACCTGGG |
| MSH2 | CCGGTCGAAAAGGCGCACTGTGG |
| MSH6 | TCCCAGGAGTACAAGAATTGAGG |

3. pegRNA information

| pegRNA | spacer sequence | 3' extension | PBS (nt) | RT T (nt) |
|---|---|---|---|---|
| HEK3_1d_ +1A ins | GGCCCAGACT GAGCACGTGA | TCTGCCATCATCGTGCTCAGTCTG | 13 | 11 |
| HEK3_2c_ 1TtoA | GGCCCAGACT GAGCACGTGA | TCTGCCATCTCGTGCTCAG | 9 | 10 |
| HEK3_2e_ 1Tdel | GGCCCAGACT GAGCACGTGA | TCTGCCATCCGTGCTCAG | 9 | 9 |
| HEK3_2e_ 1CTTins | GGCCCAGACT GAGCACGTGA | TCTGCCATCAAAGCGTGCTCAG | 9 | 13 |
| RNF2_2f_ 1GTAins | GTCATCTTAGT CATTACCTG | AACGAACACCTCAGTACGTAATGACTAAGATGA | 16 | 17 |
| HEK3_2g_ del1-5 | GGCCCAGACT GAGCACGTGA | TGGAGGAAGCAGGGCTTCCTTTCCTCTGCCGTGC TCAG | 9 | 29 |
| HEK3_2g_ del1-10 | GGCCCAGACT GAGCACGTGA | TGGAGGAAGCAGGGCTTCCTTTCCCGTGCTCAG | 9 | 24 |
| HEK3_2g_ del1-15 | GGCCCAGACT GAGCACGTGA | TGGAGGAAGCAGGGCTTCCCGTGCTCAG | 9 | 19 |
| HEK3_2g_ del1-25 | GGCCCAGACT GAGCACGTGA | TGTCCTGCGACGCCCTCTGGAGGAAGCGTGCTCA G | 9 | 26 |
| HEK3_2h_ 1His6ins | GGCCCAGACT GAGCACGTGA | TGGAGGAAGCAGGGCTTCCTTTCCTCTGCCATCA ATGATGGTGATGATGGTGCGTGCTCAG | 9 | 52 |
| HEK3_2h_ 1FLAGins | GGCCCAGACT GAGCACGTGA | TGGAGGAAGCAGGGCTTCCTTTCCTCTGCCATCA CTTATCGTCGTCATCCTTGTAATCCGTGCTCAG | 9 | 58 |

**Supplemental Figure legends**

**Figure. S1**

(A) Detailed information of gene sets used for GSEA and KEGG analysis (B) KEGG pathway analysis of eukaryotic mismatch repair from indicated gene set, highly expressed genes are colored in red while lowly expressed genes are colored in green.

**Figure. S2**

(A) Sanger sequencing result of SLC35F2 in SLC35F2 only KO (Cont), MSH6 heterozygous KO (M6 -/+) and MSH2 homozygous KO (M2 -/-) hESCs (B) mRNA expression of MSH2 and MSH6 in WT, Mock knock-out (Cont), MSH6 heterozygous knock-out (MSH6+/-) and MSH2 homozygous knock-out H9-hESCs

**Table S1**

| Normal Cells | hESCs |
|---|---|
| Skin keratinocyte HaCaT cell line | Human embryonic stem cell (HS181) |
| Neonatal foreskin keratinocyte NHEK cell line | Human embryonic stem cell (HD90) |
| Kidney epithelial cell line HEK-293 | Human embryonic stem cell (HD83) |
| Umbilical vein cell line HUVEC | Human embryonic stem cell (HS235) |
| Breast epithelial cell line HMEC | Human embryonic stem cell (HD129) |
| Testis fibroblast cell line Hs 1.Tes | Human embryonic stem cell (SA01) |
| Extravillous trophoblast cell line HTR-8_SVneo | Human embryonic stem cell (VUB01) |
| Fibroblast of skin cell line GM-5659 | Human embryonic stem cell (ES2) |
| Melanocyte cell line Hermes 2B | Human embryonic stem cell (H13) |
| Neonatal melanocyte cell line HEM-N | Human embryonic stem cell (ES4) |
| Melanocyte cell line Hermes 1 | Human embryonic stem cell (WIBR2) |
| Melanocyte cell line HEM-LP | Human embryonic stem cell (BG01) |
| Extravillous trophoblast cell line SGHPL-5 | Human embryonic stem cell (CSES4) |
| Embryonic skin fibroblast D551 cell line | Human embryonic stem cell (WIBR1) |
| Lung fibroblast cell line WI-38 | Human embryonic stem cell (H13B) |
| | Human embryonic stem cell (T3) |
| | Human embryonic stem cell (HUES6) |
| | Human embryonic stem cell (WIBR3) |
| | Human embryonic stem cell (H1) |
| | Human embryonic stem cell (H14) |
| | Human embryonic stem cell (HUES8) |
| | Human embryonic stem cell (Cythera) |
| | Human embryonic stem cell (H7) |
| | Human embryonic stem cell (H9) |
| | Human embryonic stem cell (H14A) |

**Table S1. List of hESCs and Normal cells from NextBio portal**
